# Effects of invasion level of *Prosopis juliflora* on native species diversity and regeneration in Afar region, Northeast Ethiopia

**DOI:** 10.1101/2021.03.09.434549

**Authors:** Wakshum Shiferaw

## Abstract

The study aimed to assess (i) the effects of *Prosopis juliflora* invasion on the diversity of plant species, and floristic composition at Awash Fentale and the Amibara districts of the Afar region and (ii) the effects of *P. juliflora* invasion on the regeneration potential of native woody species. Sample collection was performed in habitats of *P. juliflora* thicket, *P. juliflora* mixed with native species stands, non-invaded woodlands, and open grazing lands. A stratified random sampling technique was used for data collection. Among species of plants, the highest proportion of species, 87 (27.4%), was recorded under non-invaded woodlands, but the lowest proportion of species, 70 (22%), was recorded under open grazing lands. The invasion level of *P. juliflora* caused significantly reduced Shannon diversity index. The mean values of Shannon diversity index and species richness under *P. juliflora* mixed with native species (H’=2.22, R=14) and non-invaded woodlands (H’=2.23, R=13) were significantly higher than *P. juliflora* thicket (H’=1.96, R=12) and open grazing lands (H’=1.84, R=10). In this study, 102 trees ha^-1^native woody species were recorded under *P. juliflora* thicket, but 1252 trees ha^-1^native species were recorded under non-invaded woodlands. If the present effects of the invasion of *P. juliflora* on native species diversity were to continue coupled with a drier climate, plant diversity of the Afar flora region will be highly affected. As a result the ecosystem services will be under threat. Thus, the participation of all stakeholders and multidisciplinary research approaches should be designed for the management of invaded rangelands to reverse the situation.

## Introduction

Invasive species are either indigenous or exotic species that can heavily colonize a particular habitat [1]. A species is considered an invasive alien species when it spreads beyond its natural area of distribution [2]. *Prosopis juliflora* (Sw.) DC is among the invasive plant species native to South America, the Caribbean, and Central America [3]. *P. juliflora* has been introduced intentionally to Ethiopia particularly in the Afar region in the late 1970s for the reclamation of degraded areas in low lands [4–7). Although introduced *P. juliflora* has been providing a few uses, for instance, fuelwood and dry season fodder in rural areas, the threat posed by it in terms of invasion of fertile agricultural lands and loss of bio-diversity is looming enormously [8]. In the lowlands of Ethiopia, rangelands are subjected to different human and natural impacts. These have in turn facilitated the invasion of undesirable herbaceous weeds and woody plants in rangelands thereby threatening pastoral production systems [9]. Among woody invasions, *P. juliflora* is imposing the most jeopardy to arid and semi-arid areas in the east and northeast Ethiopia, particularly in the Afar region [10,5,11].

Land use and land cover changes, competitive ecological advantages, and climate changes are also among the key factors thought to influence the probability of invasion of *P. juliflora* [3, 12]. Use of woody species for fuelwood and construction purposes and overgrazing have also impacts on natural resources [13,14]. The seed dispersal mechanism of *P. juliflora* is also the most facilitating process for its invasiveness of the plant. Endozoochory has been recognized as the most important dispersal mechanism in the invasion *P. juliflora*, because their sugary, tasty pods attract animals and because some of their seeds remain intact after passing through some animals’ digestive systems [15].

A recent report by Shiferaw et al. [16] revealed that 1.2 million ha of land to have been invaded by *P. juliflora* constituting 12.3% of the surface area of the Afar region. Due to low moisture requirements and dispersal agents of *P. juliflora*, roadsides, river courses, farmlands, irrigation canals, wetlands, grasslands, conservation areas, and homestead areas are the most severely invaded habitats in the Afar region [17]. *P. juliflora* is dominating large areas of prime grazing land of the Afar region. Consequently, nutrient-rich palatable grasses, the main feed source for grazers are becoming progressively outcompeted [7].

There is plenty of literature showing both the negative as well as the positive environmental uses of *P. juliflora*. However, in East Africa and the Afar region of Ethiopia in particular, the problems of *P. juliflora* are outweighing positive ones in the context of ecological, socio-economic, and health aspects [5,11]. It has been noted that *P. juliflora* has a depressing effect on annual compared with perennial plants, especially on grasses in arid lands [18,19). The study by Kahi et al. [20] revealed that standing biomass, frequency, and the cover of understorey plant species to be significantly higher in the open area than under the canopies of *P. juliflora*.

Impacts of *P. juliflora* on diversity studies in the Afar region have been discussed by many researchers such as Berhanu and Tesfaye [4], [16], Kebede [21], Getachew et al. [22], and Tesfaye [23]. However, little is known about the extent of effects posed by *P. juliflora* on plant diversity, composition, and regeneration of woody species. In this study, we tried to assess the effects of *P. juliflora* invasion levels on (i) the floristic composition and diversity of plant species and (ii) the regeneration potential of native woody species in selected *P. juliflora* invaded areas of Amibara and Awash Fentale districts of Southern Afar region, Ethiopia. Then, we tested the following hypotheses: (1) native species diversity and composition are not affected by different *P. juliflora* invasion levels, and (2) regeneration patterns of woody species.

## Materials and Methods

### Description of the study area

In this study, two *P. juliflora* invaded districts from the Southern Afar region namely Awash Fentale and Amibara districts were selected for data collection. The districts were selected based on invasion of *P. juliflora* and existence of non-invaded land use and covers (*P. juliflora* invasion levels) for comparisons. Then, four sites: Diduba and Kebena from Awash Fentale; and Kurkura and Andido from Amibara district were selected. Amibara district is located in between 741-746 m asl altitudes and 9^0^19′43.83′′ N latitudes and 40^0^10′51.6′′ E longitude, whereas Awash Fentale is located at 700-1000 m asl altitude and 9^0^10′ 00′′ N and 40^0^03′33′′E (**Figure 1**).

**Figure 1.**
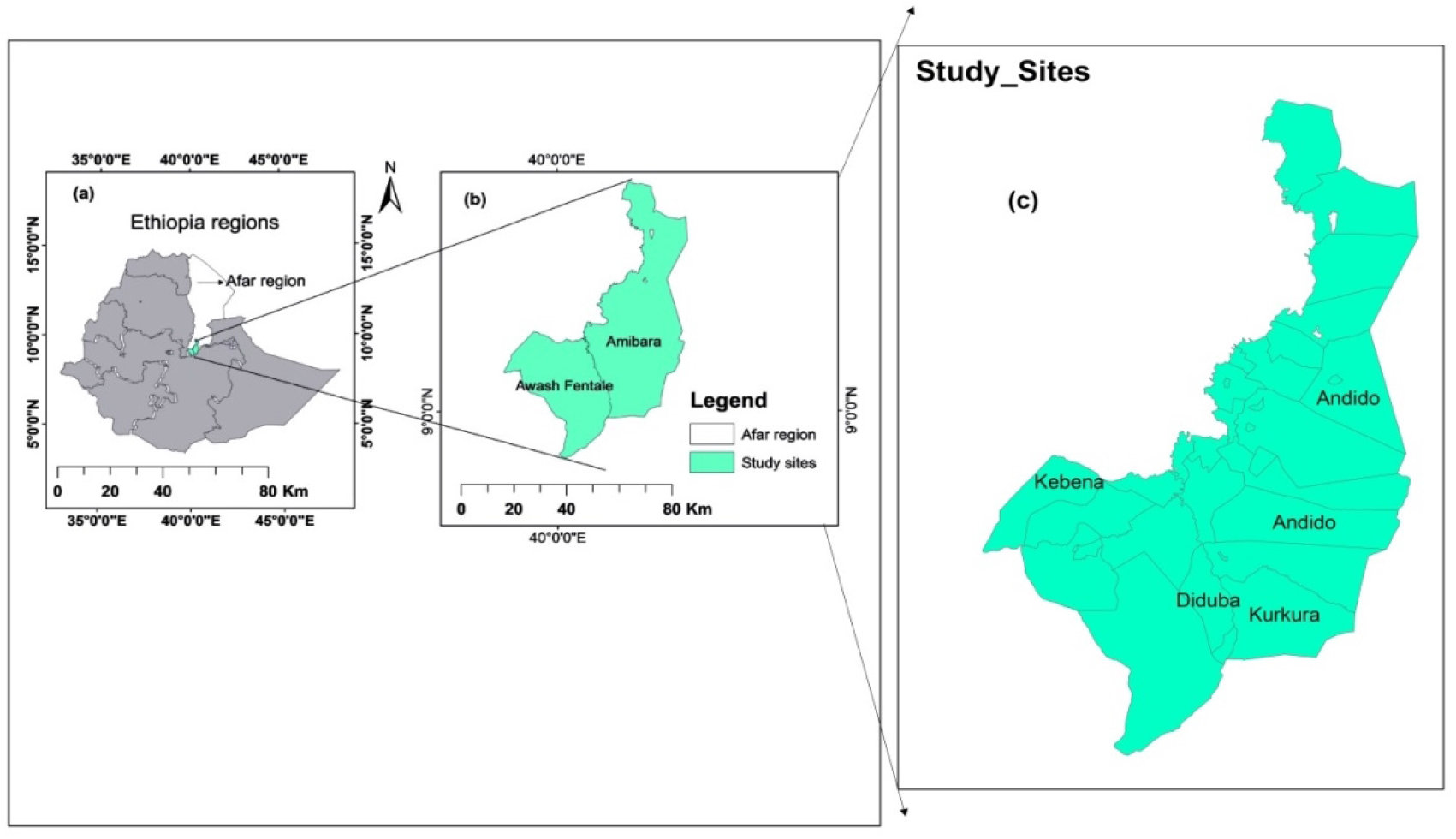
Location of the study area (Afar region) in Ethiopia (a) and location of Amibara District and Awash Fentale District in Afar region (b), and study sites (c).

The texture of the soils was usually sandy and originated from Jurassic and Cretaceous limestone and other sedimentary rocks [24]. According to FAO soil classification and ISRIC-world soil information, the soil of the Afar Floristic Region is Lithic and Eutric leptosols, and Eutric fluvisols. *Acacia-Commiphora* woodland and bushlands are among the vegetation types in Ethiopia which are characterizing the floristic region [24]. Number of population in Amibara and Awash Fentale were 83, 851, and 40,901, respectively [25]. Ninety percent of Afar people are pastoralists, while the rest are considered agro-pastoralists [26].

### Sampling design

During the preliminary reconnaissance survey, *P. juliflora* invaded patches and non-invaded lands were selected for vegetation inventory. The sites have been selected based on the severity of invasion by *P. juliflora* and the existence of non-invaded adjacent sites. Moreover, the study sites were stratified into approximately homogeneous units based on the following parameters: such as invasion levels of *P. juliflora* (invasion levels were quantified on number stems of *P. juliflora* in habitats), land use and land cover, and physiography of the sites. Thus, vegetation data were collected from (1) *P. juliflora* thicket which contained purely *P. juliflora* stems in the patch, (2) mixed *P. juliflora* with native species in the patch, and (3) non-invaded woodland and open grazing land as control which contained no *P. juliflora* were in the habitat.

Quadrants were laid at different invasion levels to collect vegetation and other environmental variables [27]. For this study, a stratified random sampling technique was used. Those quadrats for habitat categories were selected to assess the variations of *P. juliflora* invasion levels and other environmental factors on vegetation patterns, abundances, and regeneration potentials of woody species. Modified methods of habitat selection were followed: Gairola et al. [28] and Muturi et al. [29]. Thus, a total of 64 quadrats from the study sites i.e. 16 quadrats from each of four *P. juliflora* invasion levels were sampled (Plate 1).

**Plate 1.**
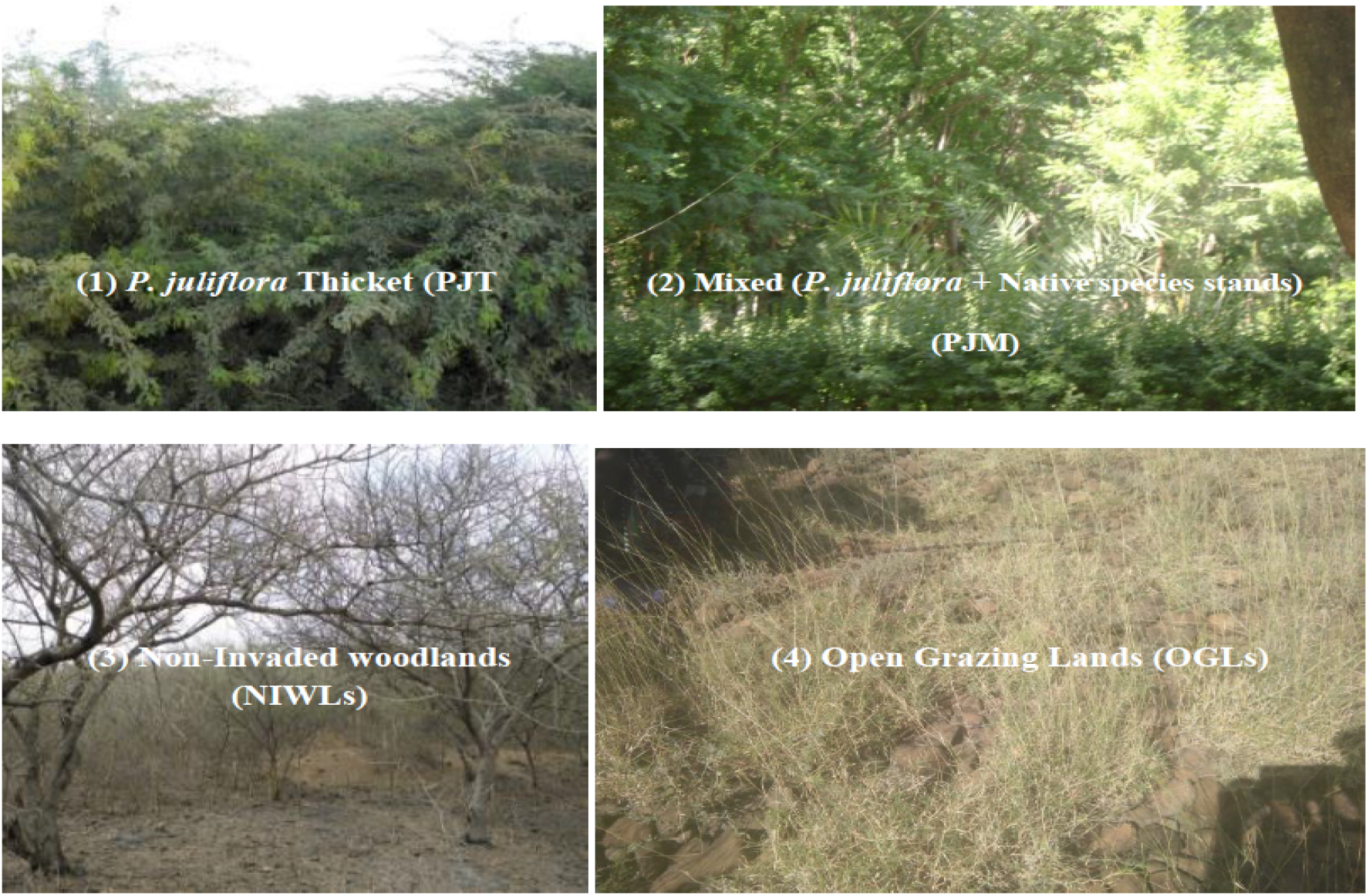
Sample plates for *P. juliflora* invasion levels/habitats in Afar Floristic Region

Vegetation data were collected under different *P. juliflora* invasion levels using quadrat sizes of 20 m x 20 m (400 m^2^) for *P. juliflora* thicket, *P. juliflora* with native species, and non-invaded woodland and open grazing lands. The first quadrat was started randomly, and then the successive quadrats were established using preferential sampling for patches and open grazing lands adjacent to each other.

### Data Collection

Individual woody categorizations were made at a height of less than 1 m and diameter at stump height (DSH) less than 1 cm for seedlings. The height 1-2 m and diameter at breast height (DBH) or diameter at stump height 1-5 cm for saplings and height greater than 2 m and DBH/DSH greater than 2 cm for tree/shrub species were measured. For regenerated seedlings (height less than 1 m), only their number was counted and recorded. In each plot, all trees or shrubs were measured and counted. Based on DBH size classes, size class structures were made for each woody species (Figure 3). Cover abundance was also identified and estimated for all growth forms using Maarel [30]. Diameter at breath height (DBH) for trees or diameter at the stump height (DSH) for shrubs was measured in the quadrats to capture all multi-stem in each individual of the shrub. Caliper and hypsometer were used to measure tree/shrub diameter and height respectively.

If a tree was branched at breast height or below, the diameter was measured separately for the branch and then averaged. Any individual with its above-ground stem growing in a cluster for woody plants (shrubs) was counted and measured as a single individual for basal area calculation following the fixed-area micro-plot method [31]. Quadrats of 1 m x 1 m (1 m^2^), 10 m x 10 m (100 m^2^), and 20 m x 20 m (400 m^2^) were used for recording seedlings/herbaceous, saplings, and tree/shrub species, respectively. That was 80 sample quadrats of 1 m x 1 m (1 m^2^), 16 quadrats of 10 m x 10 m (100 m^2^), and 16 quadrats of 20 m x 20 m (400 m^2^) were laid in each invasion levels.

The presence or absence of plant species was registered by direct counting. Percentage cover (ground cover) of herbaceous plants was estimated from the five subplots of 1 m^2^ (at four corners and one in the center of the main plot) and then the mean estimates were taken [30]. That is, all grasses and herbaceous within the marked area of the 1 m^2^ were estimated, recorded, and collected for identification. All quadrats were geo-referenced using GPS for location (coordinates) and altitudes; and a clinometer for slopes, hypsometer for the height of trees. Plant specimens were brought and stored in Addis Ababa University’s national herbarium for further identification. Plant identification and naming were followed by published flora books of Ethiopia and Eritrea.

### Data analysis

The cover/abundance values of all plant species in each plot were visually estimated using a 1-9 modified scale [27]. Evenness (*E’*) was calculated from the ratio of observed diversity to maximum diversity following Pielou [29]. Then, deviations from the means were calculated as the standard error.

The diversity of plant species were analyzed using R-software version 3.5.1 for vegetation patterns. Data series were tested for normality and homogeneity of variance. The regeneration status of woody species was determined based on the population size of seedlings, saplings, and adults. Then, all obtained vegetation patterns in terms of diversity indices and regeneration of woody species in different *P. juliflora* invasion levels were subjected to analysis of variance using the General Linear Model of SAS (version 9.0) software [30]. Significant differences among means were separated using Duncan’s multiple range tests for effects of *P. juliflora* invasion levels on vegetation patterns and regeneration potential of woody species.

## Results

### Invasive effects of P. juliflora on floristic composition and structure

In this study, 157 plant species belonging to 34 families were recorded (Appendix 1). Plant life-form distribution showed high variability among different families. In the study sites, 29 (18.5%) plant species belonged to Poaceae, 13 (8.3%) belonged to Fabaceae, and 7 (7%) species each belonged to the families Acanthaceae and Malvaceae.

Among species recorded, 87 (27.4%) were under non-invaded woodlands, but 70 (22%) were in open grazing lands. The dominant growth form was forbs under *P. juliflora* thicket, which accounted for 44 (13.8%) of all growth forms (**Figure 2**). Results also showed that 17 (5.3%) grass species were recorded under *P. juliflora* thicket, but 16 (5%) under non-invaded woodlands. Besides, 9 (2.8%) tree species were recorded under *P. juliflora* occurring with native species and 7 (2.2%) tree species in non-invaded woodlands. However, 5 tree species (1.6%) were recorded from under *P. juliflora* thicket and open grazing lands each (**Figure 2**).

**Figure 2.**
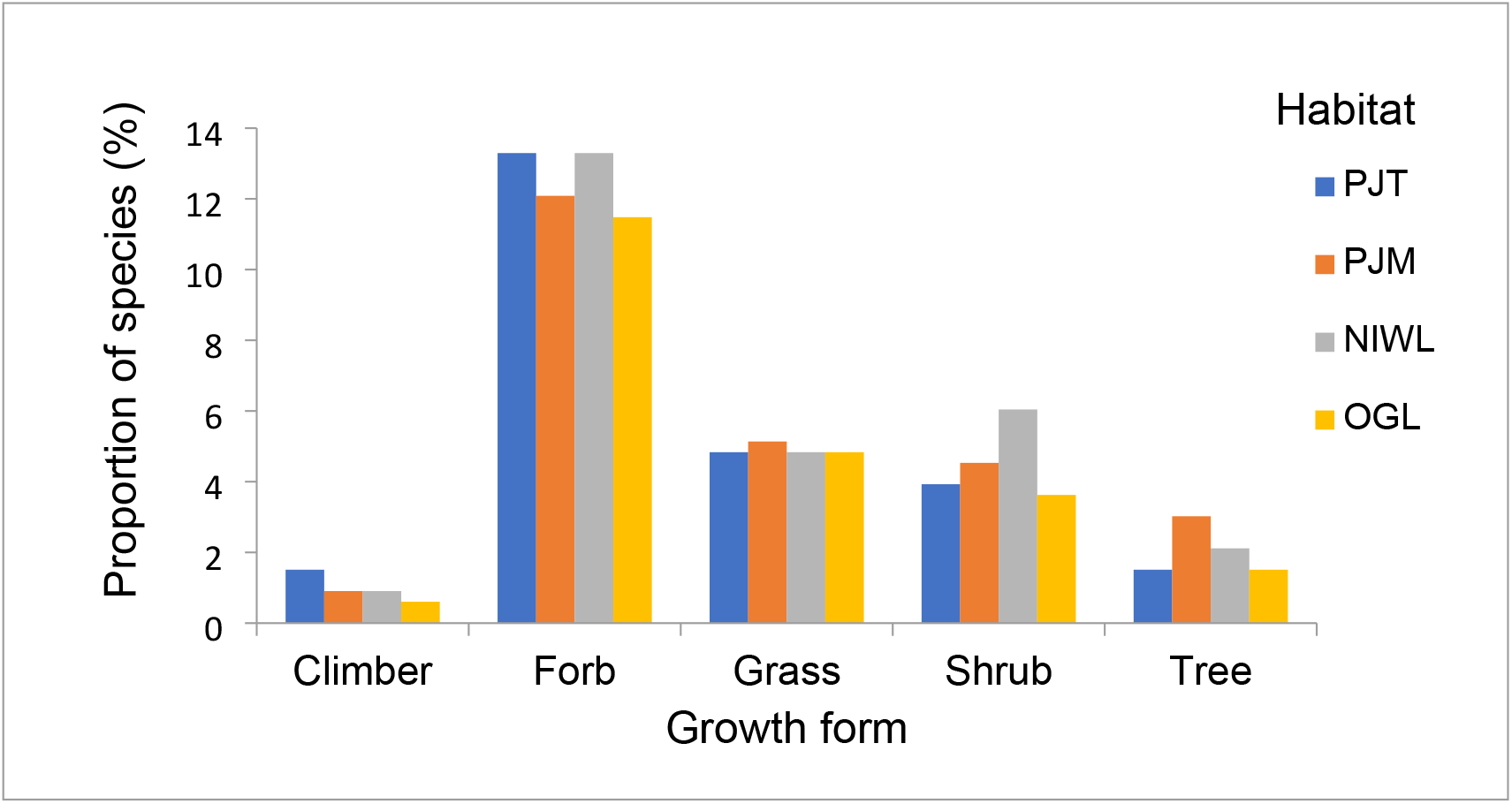
Proportion (%) of plant species in each habitat (PJT is *Prosopis juliflora* thicket, PJM is *Prosopis juliflora* mixed with naïve species, NIWL is non-invaded woodland, OGL is open grazing land).

### Invasion of P. juliflora and plant species diversity

As depicted in (Table 1), *P. juliflora* invasion levels had significantly affected the Shannon diversity index and species richness (*P <* 0.05). However, the *P. juliflora* invasion levels did not affect Shannon evenness (*P* > 0.05). On the other hand, species diversity indices (Shannon diversity index, species richness, and Shannon evenness) did not show significant difference between districts at *P* > 0.05 (Table 1).

**Table 1.**
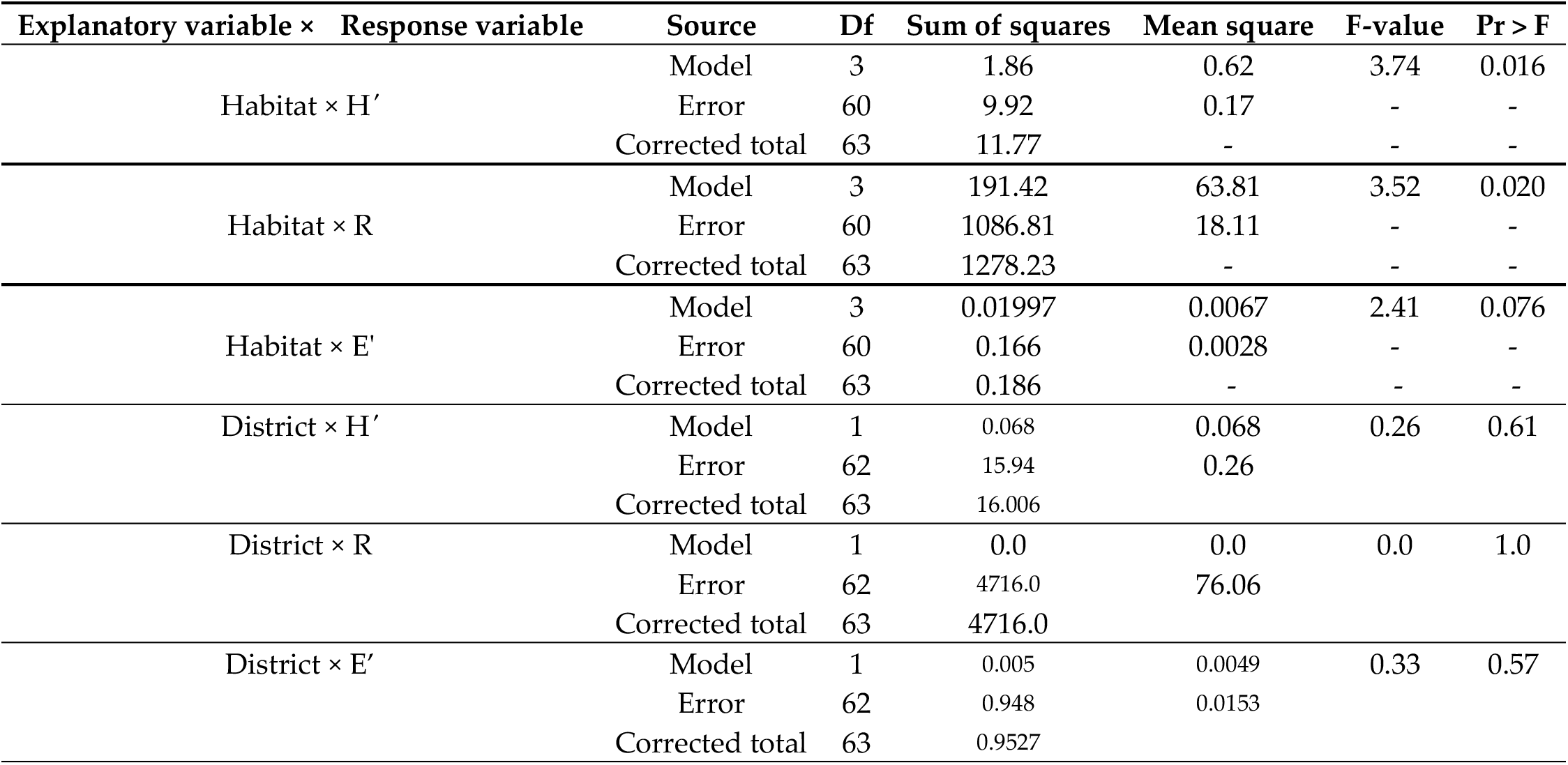
GLM showing effects location and *P. juliflora* invasion levels on species diversity at Amibara and Awash Fentale districts, Ethiopia (Significant at *p* = 0.05, H is Shannon diversity, R is species richness, E’ is Shannon evenness).

| Explanatory variable × Response variable | Source | Df | Sum of squares | Mean square | F-value | Pr > F |
| --- | --- | --- | --- | --- | --- | --- |
| Habitat × H' | Model | 3 | 1.86 | 0.62 | 3.74 | 0.016 |
|  | Error | 60 | 9.92 | 0.17 | - | - |
|  | Corrected total | 63 | 11.77 | - | - | - |
| Habitat × R | Model | 3 | 191.42 | 63.81 | 3.52 | 0.020 |
|  | Error | 60 | 1086.81 | 18.11 | - | - |
|  | Corrected total | 63 | 1278.23 | - | - | - |
| Habitat × E' | Model | 3 | 0.01997 | 0.0067 | 2.41 | 0.076 |
|  | Error | 60 | 0.166 | 0.0028 | - | - |
|  | Corrected total | 63 | 0.186 | - | - | - |
| District × H' | Model | 1 | 0.068 | 0.068 | 0.26 | 0.61 |
|  | Error | 62 | 15.94 | 0.26 | - | - |
|  | Corrected total | 63 | 16.006 | - | - | - |
| District × R | Model | 1 | 0.0 | 0.0 | 0.0 | 1.0 |
|  | Error | 62 | 4716.0 | 76.06 | - | - |
|  | Corrected total | 63 | 4716.0 | - | - | - |
| District × E' | Model | 1 | 0.005 | 0.0049 | 0.33 | 0.57 |
|  | Error | 62 | 0.948 | 0.0153 | - | - |
|  | Corrected total | 63 | 0.9527 | - | - | - |

The mean value of Shannon diversity index in Table 2 under *P. juliflora* mixed with native species (2.22) and non-invaded woodlands (2.23) were significantly higher than *P. juliflora* thicket (1.96) and open grazing lands (1.84). Moreover, the average values of species richness in Table 2 under *P. juliflora* mixed with native species (13.94) showed significantly higher than open grazing lands (9.56). Also, the Shannon evenness of non-invaded woodlands (0.87) was significantly higher than *P. juliflora* thicket (0.83) and open grazing lands (0.83) (Table 2).

**Table 2.**
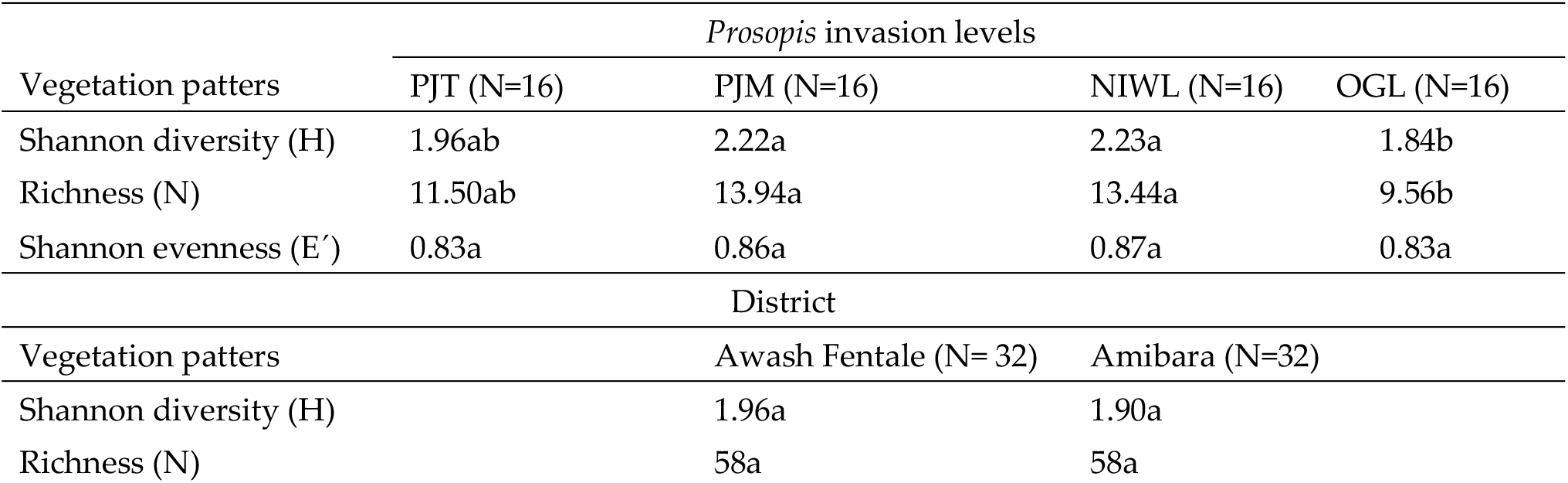
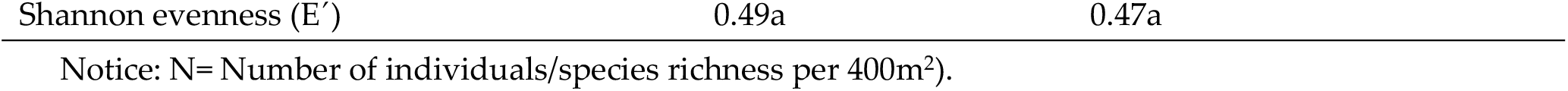
Mean values of vegetation patterns for *Prosopis* invasion levels in South Afar, Ethiopia

| Vegetation patters | Prosopis invasion levels |  |  |  |
| --- | --- | --- | --- | --- |
|  | PJT (N=16) | PJM (N=16) | NIWL (N=16) | OGL (N=16) |
| Shannon diversity (H) | 1.96ab | 2.22a | 2.23a | 1.84b |
| Richness (N) | 11.50ab | 13.94a | 13.44a | 9.56b |
| Shannon evenness (E') | 0.83a | 0.86a | 0.87a | 0.83a |
| Vegetation patters | District |  |  |  |
|  | Awash Fentale (N= 32) |  | Amibara (N=32) |  |
| Shannon diversity (H) | 1.96a |  | 1.90a |  |
| Richness (N) | 58a |  | 58a |  |

Notice: N= Number of individuals/species richness per 400m<sup>2</sup>).

**Table 3.** General Linear Model showing the effects of *P. juliflora* invasion levels on regeneration status of woody species at Amibara and Awash Fentale districts, Ethiopia.

| Response variables | Source | Df | Sum of squares | Mean square | F-value | Pr > F |
| --- | --- | --- | --- | --- | --- | --- |
| Habitat × Trees | Model | 2 | 36907.7 | 18454 | 0.94 | 0.41 |
|  | Error | 21 | 410318.9 | 19539 | - | - |
|  | Corrected total | 23 | 447226.7 |  |  |  |
| Habitat × Saplings | Model | 2 | 32.66 | 16.33 | 0.71 | 0.49 |
|  | Error | 1592 | 36640.01 | 23.01 | - | - |
|  | Corrected total | 1594 | 36672.67 | - | - | - |
| Habitat × Seedlings | Model | 2 | 69.28 | 34.64 | 1.06 | 0.35 |
|  | Error | 1592 | 51796.23 | 32.53 | - | - |
|  | Corrected total | 1594 | 51865.51 | - | - | - |

### Invasion of P. juliflora and size class structures of woody species

As can be seen from the result in (Table 4) tree, saplings and seedlings are shown to be statistically not affected by the invasion of *P. juliflora* at the different invaded habitats (*P >* 0.05). The lowest total densities of trees were recorded under *P. juliflora* thicket (102 stems/ha). But, the highest total densities of trees were recorded under non-invaded woodlands (1252 stems/ha). Moreover, the highest total density of seedlings was recorded under *P. juliflora* mixed with native species (358 stems/ha). However, the lowest total densities of seedlings were recorded under *P. juliflora* thickets (153 stems/ha) (Table 4).

**Table 4.** Mean values of the regeneration status for *P. juliflora* invaded and non-invaded habitats at Amibara and Awash Fentale districts, Ethiopia (PJT is *P. juliflora* thicket, PJM is *P. juliflora* mixed with native species, NIWL is non-invaded woodlands, stems/ha).

| Growth Stages | Habitat |  |  |
| --- | --- | --- | --- |
|  | PJT | PJM | NIWL |
| Trees | 102a | 585b | 1252b |
| Saplings | 151a | 334b | 324c |
| Seedlings | 153a | 358b | 242c |

### Invasion of P. juliflora and size class structures of a selected population of woody species

Results showed that the pattern of population structure for a given species can be roughly grouped in one of four basic types: Type I (inverted J-shape), Type II (bell-shaped type), Type III (J– shape), and Type IV (U-shaped). Type I is a pattern in which a diameter size class distribution displays a greater number of smaller seedlings than large ones. Type II is characteristic of species that show discontinuous, irregular, and or periodic recruitment. Type III reflects a species whose regeneration is severely limited. Type IV is characteristic of species that show a lower number of saplings or mid-class distribution than seedlings and trees.

The results in Figure 3 (3b&f) showed that the population of *P. juliflora* and *Acacia tortilis* (Frossk.) Hayne both under *P. juliflora* mixed with native species, their regeneration patterns had U-shaped (Type IV) type. Meanwhile, the population of *Cordia sinensis* Lam under non-invaded woodlands; *Cadaba rotundifolia* Forssk and *Grewia tenax* (Forssk.) Fiori under *P. juliflora* mixed with native species showed that regeneration patterns bell-shaped type (Type II) (Figure 3i&j). Furthermore, in Figure (3d&j) the population of *C. rotundifolia*, under non-invaded woodlands, and *Salvadora persica* L. under *P. juliflora* mixed with native species were showed the regeneration patterns of inverted J-shape (Type I). On the other hand, the population of *A. mellifera, S. persica* and *A. tortlis* under NIWL; and *P. juliflora, G. tenax, A. oerfota, A. senegal, A. mellifera* and *C. rotundifolia* under *P. juliflora* thickets; and *A. nilotica* under non-invaded wood lands and woody species like *A. mellifera, A. oerfota, A. tortilis, G. tenax, S. persica, C. rotundifolia* under open grazing lands resembled the limited regeneration patterns of J–shape (Type III) in the study areas (Figure 3, b, d, g, h, i & j).

**Figure 3.**
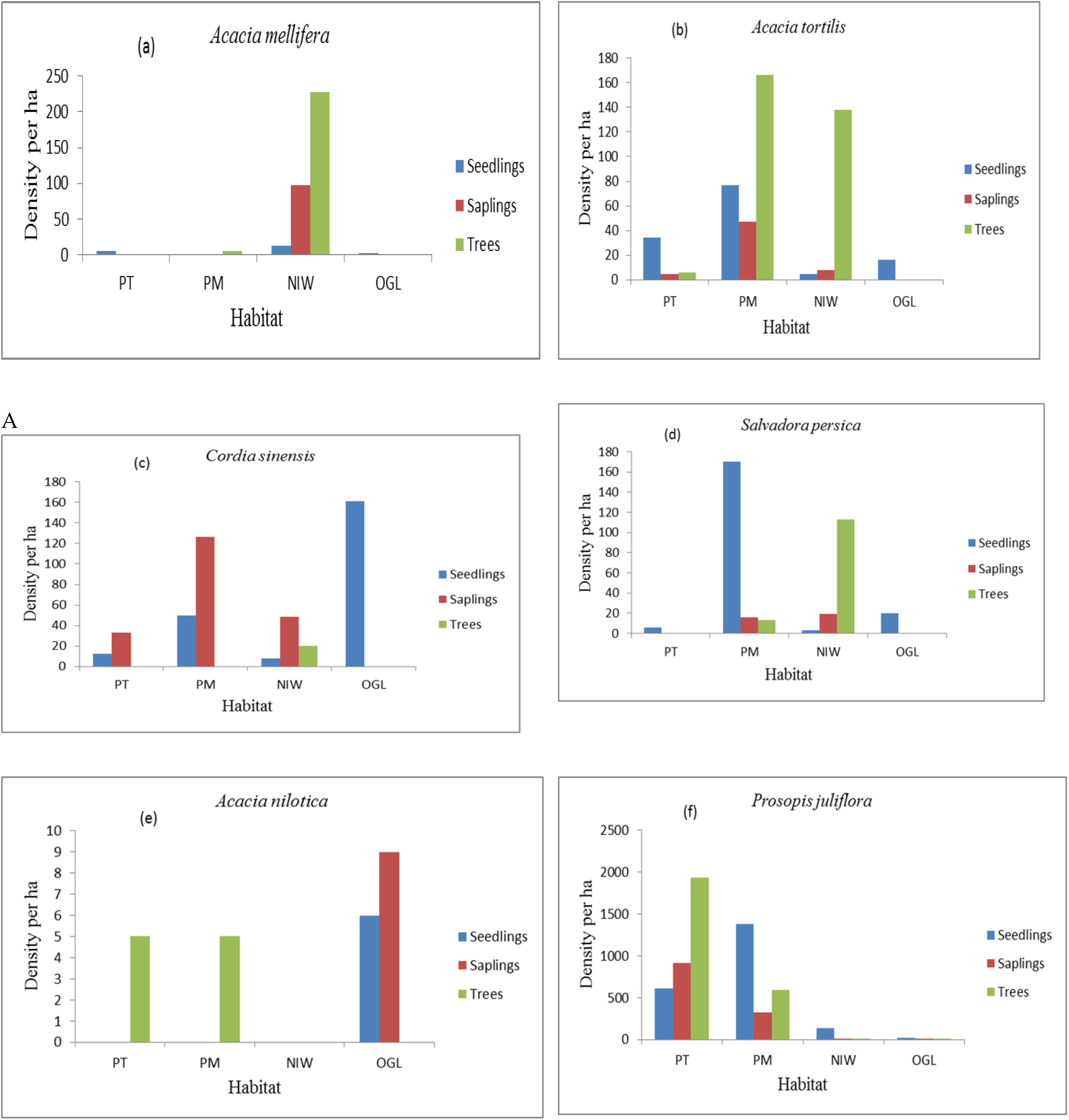
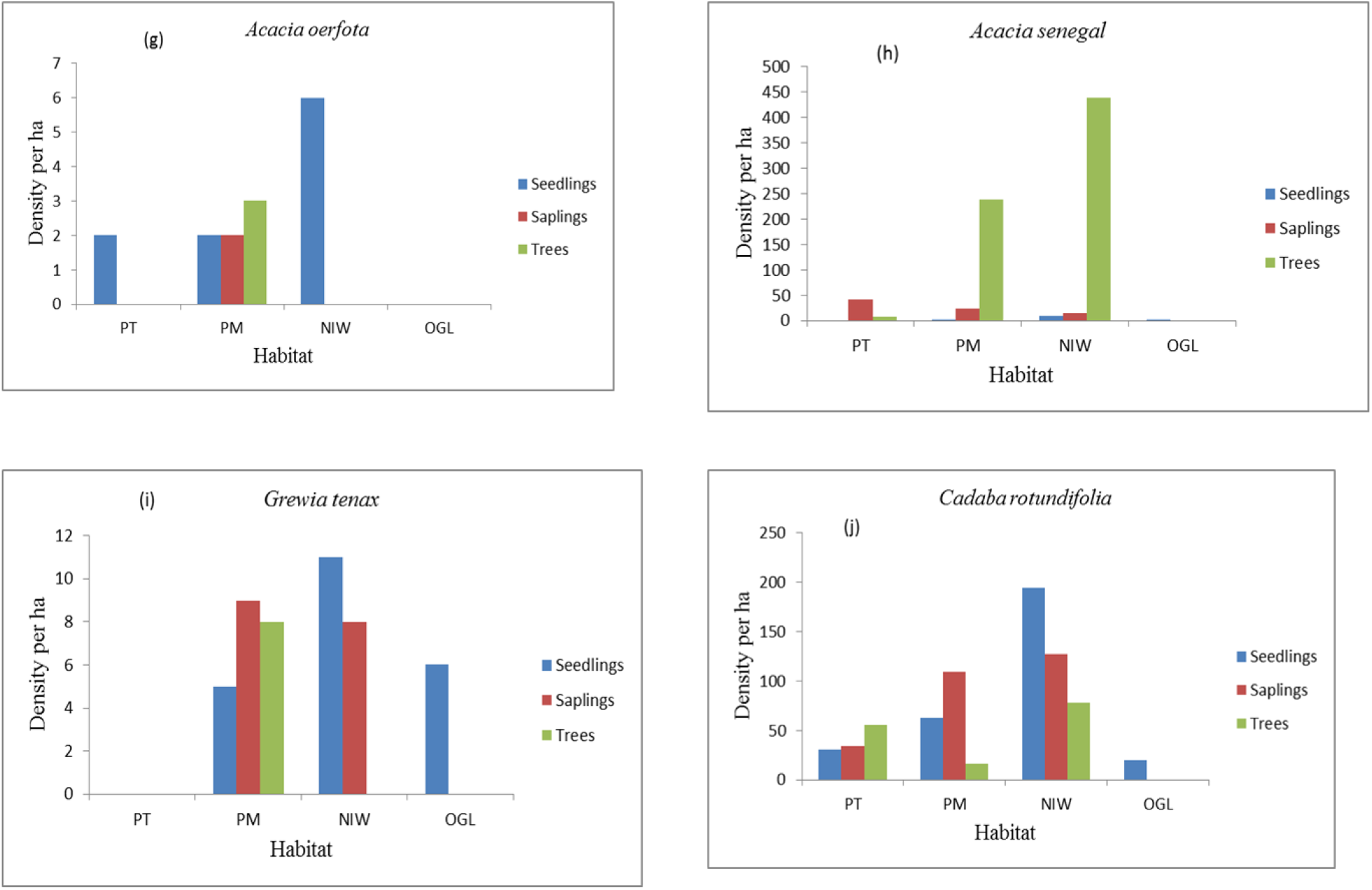
a-j. Size class structures of ten species population in each habitat (PT = ***P. juliflora*** thicket stands, PM= ***P. juliflora*** with native species stands, NIW = non-invaded woodland, OGL = open grazing land).

## Discussion

### P. juliflora and floristic composition

In this study, the number of families identified was comparable with research reported by Aggarwal [33] from the pre-urban region of India and findings by Demissie [34] in Awash National Park in Ethiopia. Other findings by Mukherjee et al. [32] showed that both the number of species and families were lower than the number of families recorded in the present study.

The species richness under the canopy of *P. juliflora*, in this study, was lower compared with those reported by Patel et al. [36] for Delhi, Gujarat, and Rajasthan in India. Our study also revealed that the number of species under *P. juliflora* thicket was higher than that of the number of species under the *P. juliflora* thicket on Delhi ridge of India [37]. Similar trends on the number of associated species were also observed in sites with a lower density of *P. juliflora* under *P. juliflora* with native species and outside (non-invaded woodlands) canopies in UAE [38].

The presence of more climbers under *P. juliflora* thicket compared to other habitats might be due to the presence of woody species that help the climbers as support. The trees in the habitats also serve as safety factors to resist a range of natural factors, particularly high winds [39].

A higher number of forb species were also observed under *P. juliflora* thicket than other habitats. The availability of forbs contributes to suitable resource islands which had out-competed other species under the thicket of *P. juliflora* canopies [40,41]. On the other hand, grass species were higher under *P. juliflora* with native species than other habitats. In these particular cases, *P. juliflora* invasion might affect the growth of grasses due to the inhibition of light penetration and allelopathic chemicals under the thicket of *P. juliflora* [18]. In the present study, the decline in tree forms under the habitats of non-invaded woodlands could be attributed to anthropogenic impacts (i.e., exploitation of trees for firewood and construction) and browsing [13,42,43].

### P. juliflora invasion and species diversity

Species diversity has been recognized as an important component of the sustainable development of *P. juliflora* with native species [26]. This is because of ecosystems with diverse species resilient for the competition of resources with *P. juliflora*. Species diversity was lower both under open grazing lands (1.84) and *P. juliflora* thicket (1.96) compared to *P. juliflora* mixed with native species (2.22) and non-invaded woodlands (2.23) habitats. The reason might be disturbance intensities under open grazing lands and effects of canopy *P. juliflora* which had dominated other native species. The species diversity under non-invaded woodlands and *P. juliflora* mixed with native species stands were comparable to the report made by Zerga [44] in Awash National Park of Ethiopia and by Aggarwal, [33] at Peri-urban Region in India.

Other findings such as those of Kumar and Mathur [45] indicate higher species diversity than that of the present study. The reason might due to grazing impacts and moisture stress in the study areas. In the present study, the average values of species richness ranged from open grazing lands (9.6) to *P. juliflora* with native species (13.9). The higher the species richness under *P. juliflora* with native species and non-invaded woodlands contributes to anthropogenic impacts, grazing, and disturbance intensities, and the decline of *P. juliflora* invasions and its canopy effects which *P. juliflora* is an aggressive invader, canopy effects were consistently and strongly negative on species richness [46]. In this study, the mean values of species richness were similar under *P. juliflora* with native species to *P. juliflora* invaded forest and arid grazing lands at the Gujarat state of India, but the mean values of species richness in our study were higher under *P. juliflora* with native species and *P. juliflora* thicket habitat than *P. juliflora* invaded areas of Reserve Vidis and open grazing of arid grazing lands at Gujarat state of India [45]. In all of the habitats of this study on the contrary, we found lower species richness than Aggarwal [33] in a Peri-urban Region of India during rainy, winter and, summer seasons.

### P. juliflora invasion and size class structures of woody species

Quantitative analysis of the regeneration status of woody species recorded in this study may provide baseline information to design and formulate conservation and management strategies for *P. juliflora* dominated woodlands. The regeneration status of woody species of any vegetation is determined based on densities of seedlings and saplings [47]. The ratio of various age groups in a population determines the reproductive status of the population and indicates the future course [48].

In this study, the highest species diversity of trees, saplings, and seedlings were recorded under *P. juliflora* thicket, *P. juliflora* with native species stands, and under non-invaded woodlands in that order. The decline in the number of trees under non-invaded woodlands is accounted for by the effects of human impacts, livestock grazing, and disturbance intensities [13]. Besides, the increase in the density of individual stems in the form of the tree and seedling stages under non-invaded woodlands and *P. juliflora* mixed with native species stands might be due to its allopathic substances that would inhibit the growth of associated native species [3,49,22].

Due to illegal cutting, the abundance of tree species under non-invaded woodlands and *P. juliflora* with native species stands were similar to the research findings by Patel et al. (2012) in western Kachchousehold of Gujarat in India. However, our findings were contrary to results reported by Muturi et al. [29] in which the distribution of trees under *P. juliflora* with native species stand was higher than non-invaded woodlands and *P. juliflora* thicket in the Turkwel riverine forest of Kenya. The reason could be the variations in the management of vegetation types and age of *P. juliflora* variations which might not also affect native species under the *P. juliflora* with native species stands in Kenya.

### P. juliflora invasion levels and size class structures of selected woody species

Regeneration is a crucial phase of vegetation management as it maintains the desired species composition and stocking and can be predicted by the structure of the population [50]. Different categories of regeneration status were designated following [51,50].

*A. mellifera* under *P. juliflora* thicket and *P. juliflora* mixed with native species stands, *A. oerfota* under *P. juliflora* thicket and non-invaded wood lands, *G. tenax, A. senegal, A. nilotica* and *C. rotundifolia* under open grazing lands, and *P. juliflora* under non-invaded woodlands and open grazing lands, and *G. tenax* under non-invaded woodlands showed new regeneration categories. Whereas *C. rotundifolia* and *P. juliflora* under *P. juliflora* thicket, *A. mellifera* and *A. senegal* under *P. juliflora* mixed with native species stands, and *A. senegal, S. persica* under non-invaded woodlands, *C. rotundifolia* and *G. tenax* under *P. juliflora* thicket showed fair regeneration categories.

Furthermore, *S. persica* and *C. rotundifolia* under non-invaded woodlands showed good regeneration in the study areas. In contrast to our findings, the findings of Endris et al. [52] showed that *P. juliflora* under *P. juliflora* mixed with native species stands and *A. mellifera* under non-invaded woodlands showed good regeneration profiles in the Hallideghie wildlife reserve, Northeast Ethiopia. Furthermore, the density of seedlings for *P. juliflora* and *A. tortilis* under *P. juliflora* mixed with native species stands, non-invaded woodlands, and *P. juliflora* thicket also showed inconsistent patterns of densities in comparison to reports by Muturi et al. [29] that show higher densities in comparison to trees and saplings.

## Conclusions

It has been demonstrated in this study that species diversity decreased under grazed lands and *P. juliflora* thicket. *P. juliflora* invasion thus remains a threat to the overall native species diversity and eventually harming the rangelands and livestock production. Under *Prosopis juliflora* thicket, higher DBH classes of native woody species were dominated by *P. juliflora*. But, lower DBH size classes (seedlings) were recorded under *P. juliflora* mixed with native species stands implied that the effects of *P. juliflora* on the individuals declined. Thus, appropriate silvicultural techniques (e.g. thinning) of *P. juliflora* should be practiced to lessen the invasiveness of the species. Further studies about the phenology of *P. juliflora* are vital to know the seasons of seed dispersal to manage its invasiveness. Furthermore, the regional natural resource office should provide alternative energy sources such as solar radiation and *P. juliflora* based biogas plants to alleviate the devastation of other indigenous woody species in the region. Thus, long-term effects of *P. juliflora* species on soil seed bank, soil properties, the progress of *P. juliflora* on environmental threats, and its socioeconomic impacts on other invaded areas should also be investigated in the future.

## Author Contributions

All authors have equally contributed to the manuscript

## Acknowledgments

The authors are thankful to Arba Minch and Addis Ababa Universities for financing the project. The first author special thanks Afar pastoral communities for their cooperation and assistance during data collection. Department of natural resources management in pastoral and agro-pastoral districts of the Amibara and Awash Fentale districts are acknowledged for material and human resources assistance during site selection and data collection. All the members of the National Herbarium of Addis Ababa University are also appreciated for their facilitation of materials in the herbarium for plant identification.

